## Supplementary Informations for "Neurovascular coupling and briefCO_2_ interrogate distinct vascular regulations"

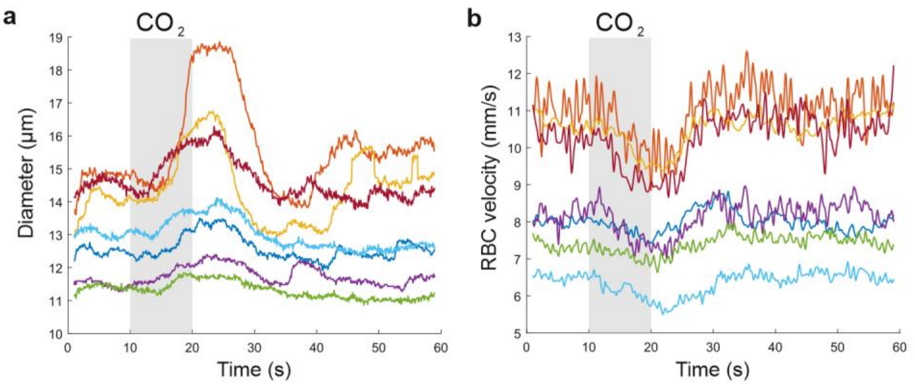


**Supplementary Figure 1. Raw data from Figure 2.**
**a,** Diameter and **b,** red blood cell (RBC) velocity simultaneously measured in pial arterioles using broken line scans to calculate blood flow changes upon brief$CO_2$ stimulation. n = 7 vessels, 6 mice.

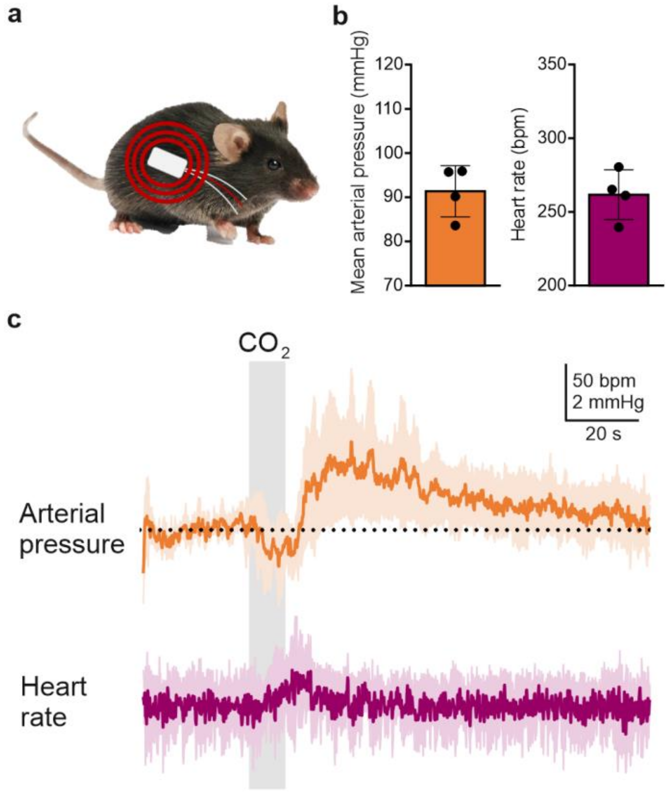


**Supplementary Figure 2. brief$CO_2$ causes an early drop in blood pressure.**
**a,** Mice were chronically implanted with a blood pressure telemetric system as described in Methods. **b,** Resting mean arterial pressure and heart rate under dexmedetomidine sedation. Note that dexmedetomidine sedation leads to a marked reduction of heart rate and a mild hypotension (expected values in awake C57/Bl6 mice during the diurnal phase[1,2]: 550 bpm, 100 mmHg). **c,** Top, Brief$CO_2$ induces a rapid drop in arterial pressure of about 1 mmHg, lasting the duration of the stimulus (10 s), and followed by a positive rebound. Bottom, Brief$CO_2$ induces a reversible increase in heart rate (4-8 trials, 4 mice). Data are represented as mean ± SD.

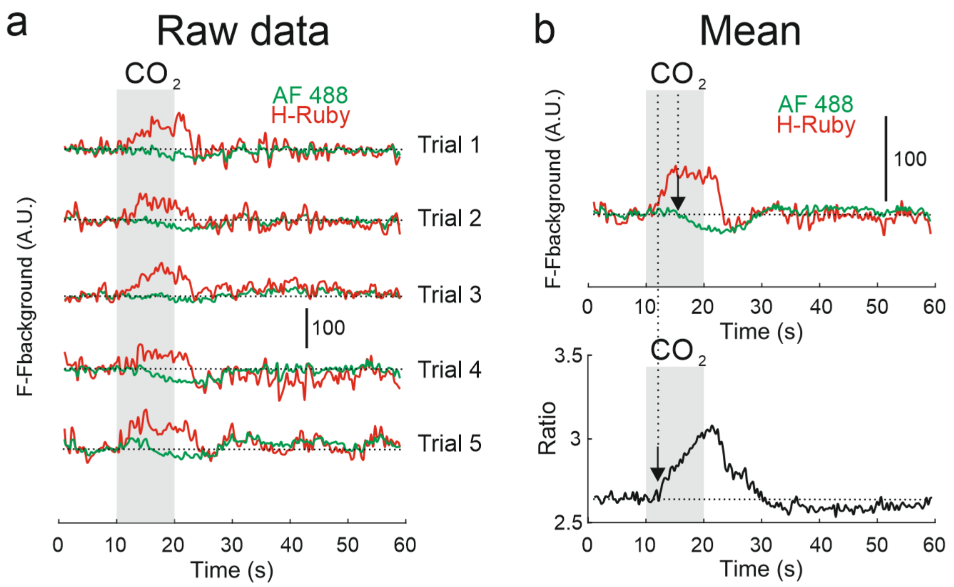


**Supplementary Figure 3. The necessity of dye ratioing.**

**a,** AF 488 (pH insensitive) and H-Ruby (pH sensitive) fluorescence changes measured in a pial arteriole in response to 5 consecutive brief$CO_2$ stimulations. For both dyes, $F_{background}$ was measured in the tissue outside the pial vessel. **b,** Top, averages of the 5 trials. Note the rapid increase of the H-Ruby signal, indicating the early onset of pial arteriole acidosis, and the delayed drop of AF488 fluorescence, in phase with the vessel dilation and reporting light absorption by hemoglobin. This shows the importance of dual simultaneous measurements to get rid of the bias due changes in absorption of excitation light by hemoglobin. Bottom, the fluorescence ratio reveals the real shape the pH change.

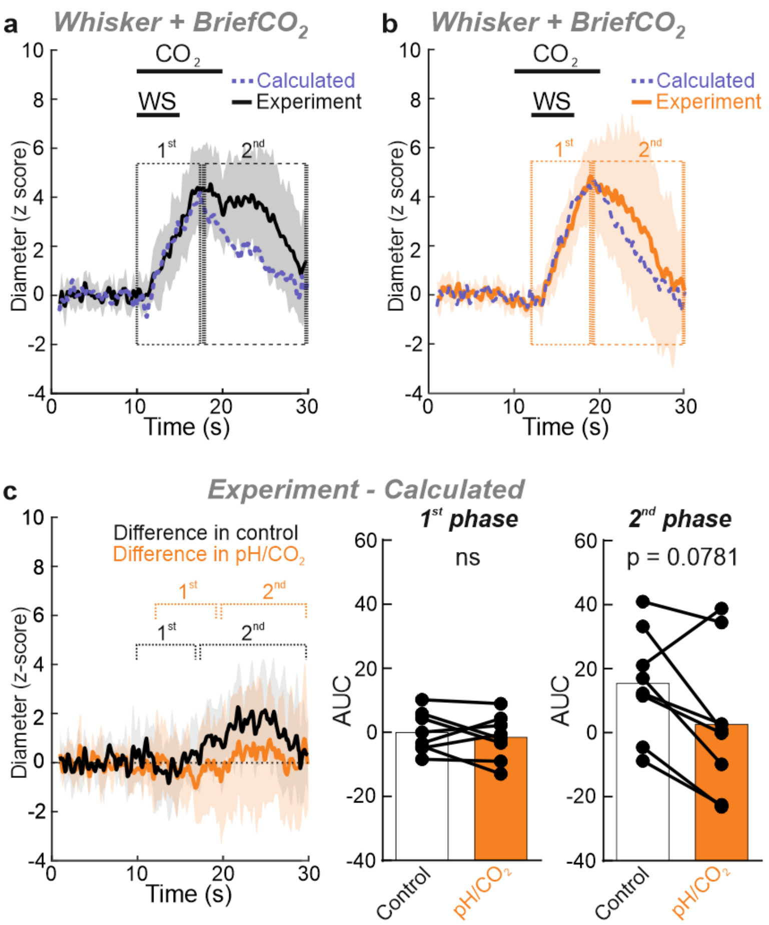


**Supplementary Figure 4. Additivity of NVC and responses to brief$CO_2$.**

**a,** Calculated summation (dotted purple line) and experimental data (black solid line) of the responses to whisker (5 Hz, 5 s) stimulation and brief$CO_2$ (20% $CO_2$, 10 s) applied at the same time. **b,** Calculated summation (dotted purple line) and experimental data (orange solid line) when whisker stimulation is delayed by 2 s from the onset of brief$CO_2$. **c,** Left, Difference between calculated and experimental vascular responses in both conditions. Middle, AUC for area under the curve during the first phase of the response (between 10-17 s and 12-19 s, i.e. when WS is simultaneous or delayed by 2s from brief$CO_2$) (Paired data, Wilcoxon sum rank test, ns, $p = 0.4609$. Right, AUC during the second phase of the response. Paired data, Wilcoxon sum rank test, ns, $p = 0.0781$. $n = 8$ vessels, 5 mice). Data are represented as mean ± SD.

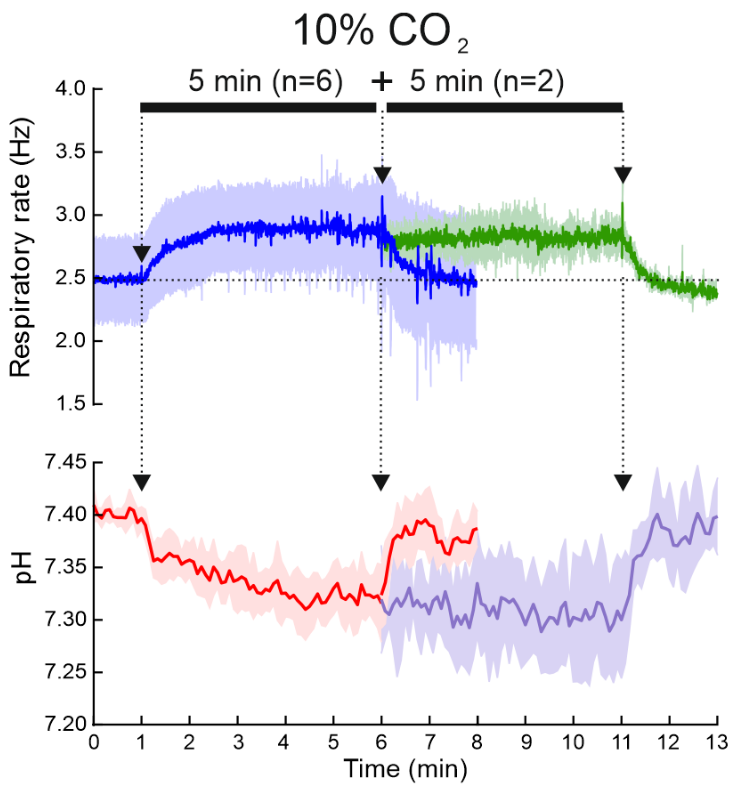


**Supplementary Figure 5. Prolonged $CO_2$ stimulation generates a sustained blood acidosis.**

10% $CO_2$ stimulation for 5 minutes (1$^{st}$ segments, blue and red traces, n = 6 experiments, 3 mice) or 10 minutes (2$^{nd}$ segments, green and purple traces, n = 2 additional experiments, 2 mice) induces a sustained increase of respiratory rate (top) and a continuous arteriolar acidification (bottom). Data are represented as mean ± SD.

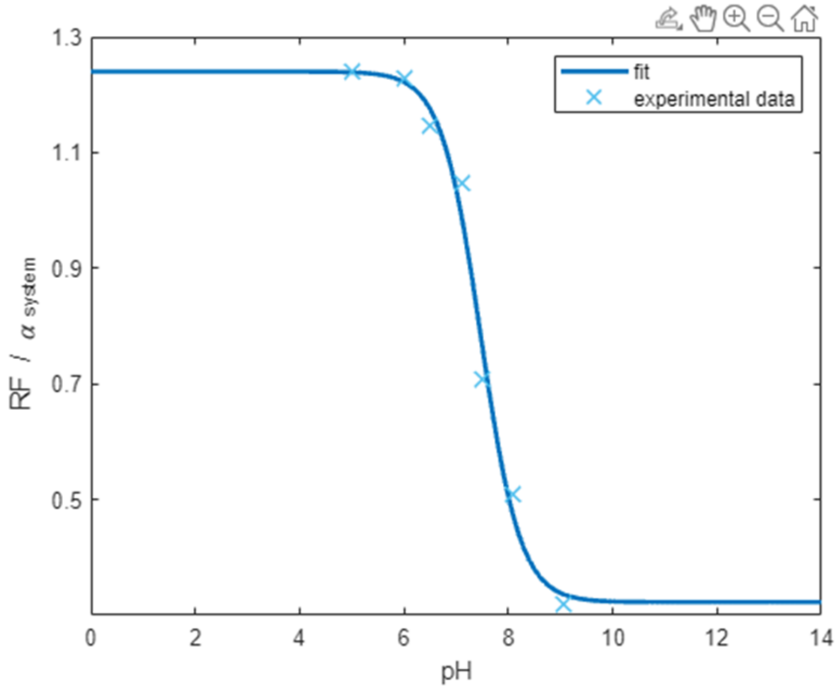


**Supplementary Figure 6. Calibration used for the look-up table.**
A look-up table was established by simultaneously measuring red and green fluorescence of a solution of H-Ruby-dextran and AF488 dextran in fresh rat blood plasma. pH was controlled with a pH sensitive electrode. The ratio of the average red fluorescence over the average green fluorescence (RF) was normalized by the relative sensitivity of the system ($\alpha_{system}$) and plot as a function of pH, fitted with a sigmoid function and used as a reference to calibrate in vivo pH measurements.

| | |
|---|---|
| Diameter | 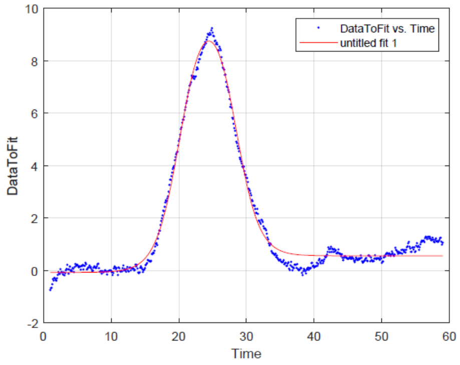 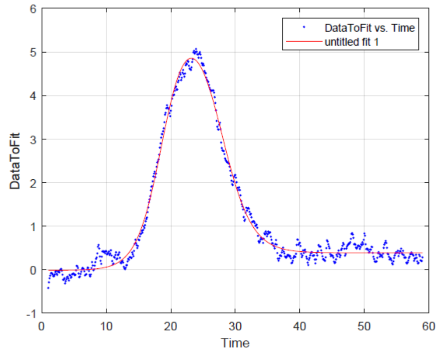<br>Average diameter responses of pial (left) and penetrating arterioles (right) (z score, data from Figure 1) were fitted with the sum of two sigmoids: (a0/(1+(exp(-(x-x0)/T0)))+a1/(1+(exp((x-x1)/T1)))+c) |
| Respiratory rate | 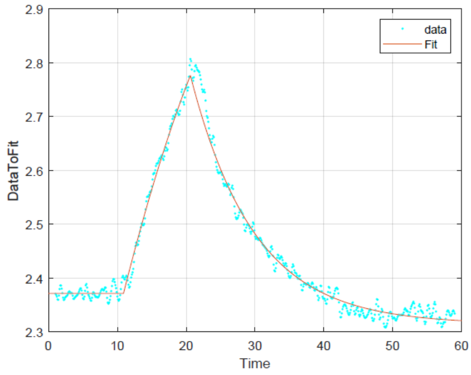 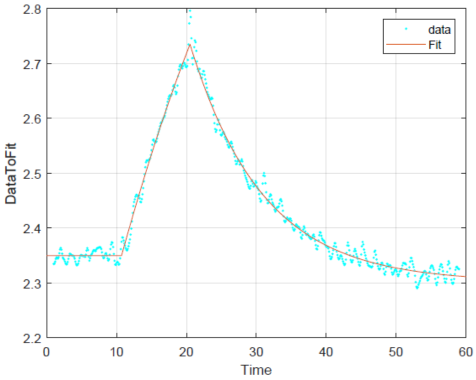<br>Average respiratory rate responses (measurements paired with pial (left) and penetrating arterioles (right); data in Hz, data from Figure 1) were fitted with a model function, composite of a steady state baseline, a second-order polynomial for the rising phase (a1*(x-T1)^2 + b1*(x-T1) + c1), and the sum of two exponentials for the decay phase: a2/(1+(exp(-(x-x2)/T2))) + a3/(1+(exp((x-x3)/T3))) + c2. |
| RBC velocity (pial arterioles) | 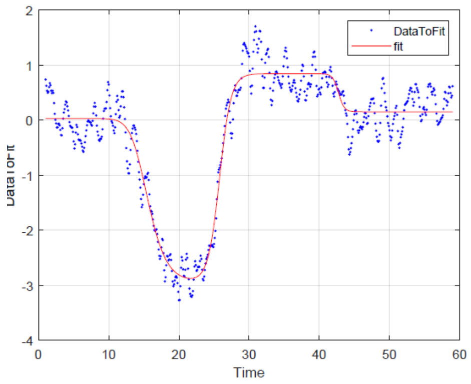<br>Average velocity responses of pial arterioles (z score, data from Figure 2) were fitted with a sum of two sigmoids: and the square of a 3rd sigmoid: |

| | |
|---|---|
| | a0/(1+(exp(-(x-x0)/T0)))+a1/(1+(exp((x-x1)/T1)))+a2/(1+(exp(-(x-x2)/T2)))^2+c. |
| GCaMP6/8 fluorescence | 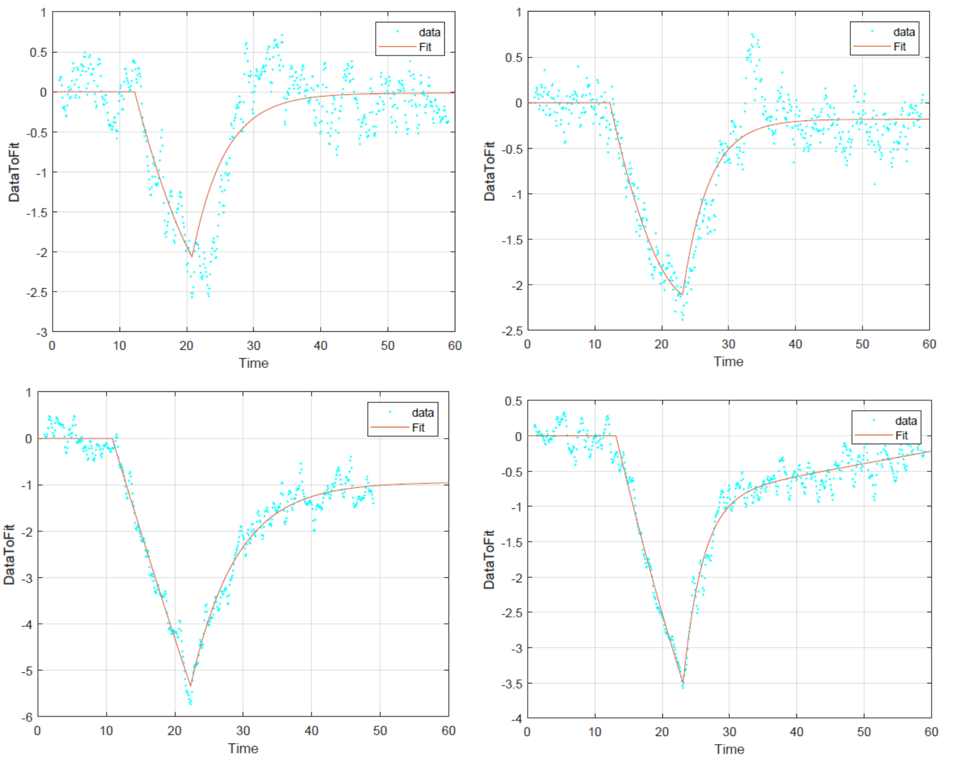<br>Average fluorescence responses of smooth muscle cells (top left), endothelial cells (top right), astrocytes (bottom left), and neuropil (bottom right) (z score, data from Figure 3) were fitted with a model function, composite of a steady state baseline, a second-order polynomial for the initial phase of the response (a1*(x-T1)^2 + b1*(x-T1) + c1), and the sum of two exponentials for the decay phase: a2/(1+(exp(-(x-x2)/T2))) + a3/(1+(exp((x-x3)/T3))) + c2. |
| CBV (fUS) | 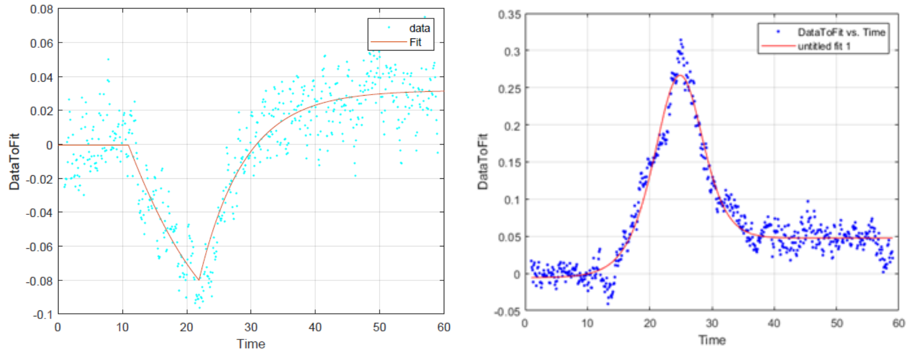<br>Average CBV responses in the carotid artery (left, z score, data from Figure 2) were fitted with a composite function made of a steady state baseline, a second-order polynomial for the initial phase of the response (a1*(x-T1)^2 |

| | |
|---|---|
| | + b1*(x-T1) + c1), and an exponential for the decay phase : a2/(1+(exp(-(x-x2)/T2))) + c2 .<br><br>Average CBV responses in the cortex (right, z score, data from Figures 2 and 5) were fitted with the sum of two sigmoids: (a0/(1+(exp(-(x-x0)/T0)))+a1/(1+(exp((x-x1)/T1)))+c). |
| Blood flow in arterioles | 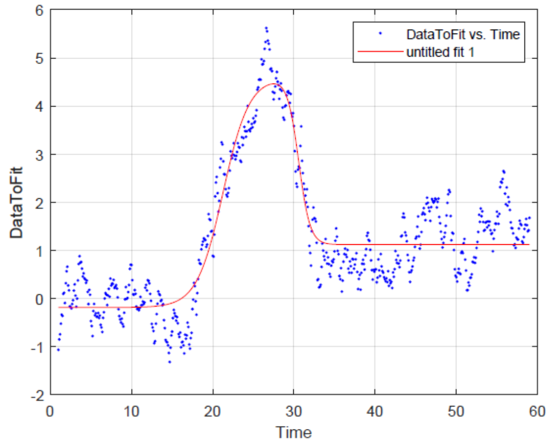<br>Average blood flow responses in the pial arterioles (z score, data from Figure 2) were fitted with the sum of two sigmoids: (a0/(1+(exp(-(x-x0)/T0)))+a1/(1+(exp((x-x1)/T1)))+c). Note that the negativity was on purpose not considered for the fit. |
| Intravascular pH | 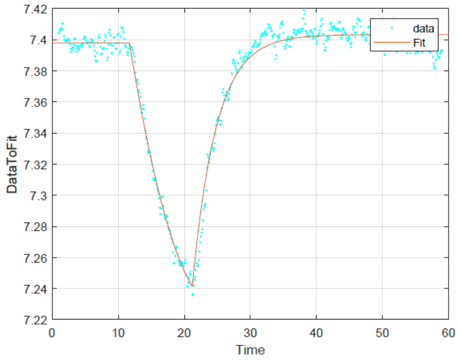<br>Average intravascular pH (measured from the ratio of H-Ruby-dextran and AF488-dextran) was fitted with the same function as for GCaMP fluorescence. |

**Supplementary Table 1. Average responses, fits (in red) and model functions used to compute the fits.**

### Synthesis of H-Ruby Dextran conjugate

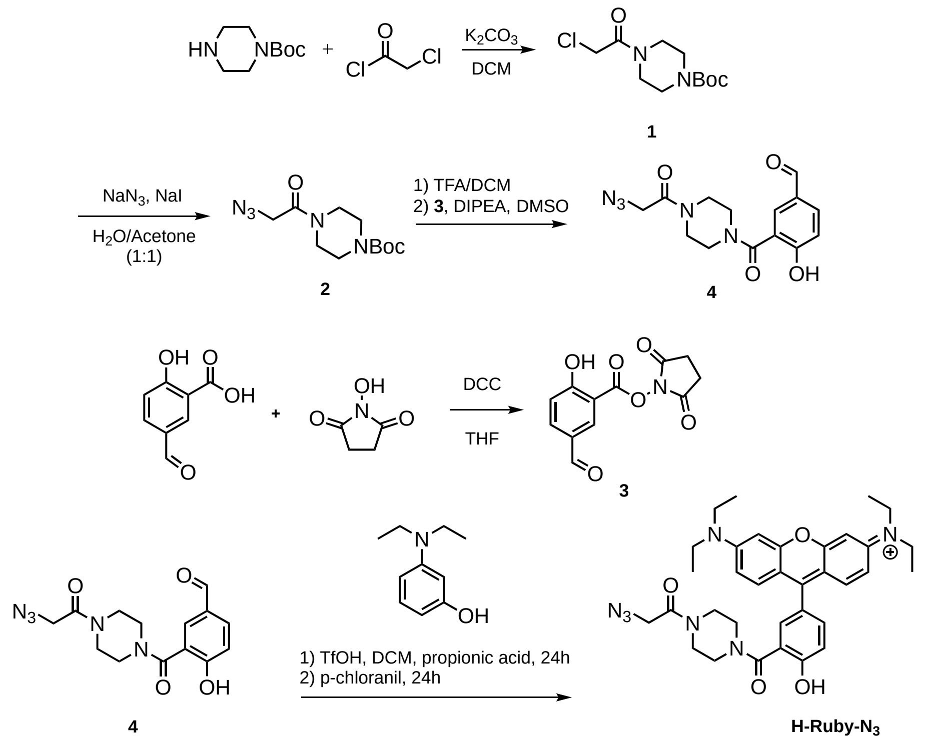


#### Synthesis of H-Ruby-$N_3$

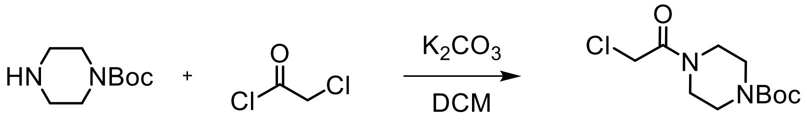


**1**

**1.** To a solution of 1-Boc-piperazine (1.0 g, 5.4 mmol, 1 eq.) and $K_2CO_3$ (0.89 g, 6.4 mmol, 1.2 eq.) in DCM (10 mL) was added chloroacetyl chloride (470 µL, 5.9 mmol, 1.1 eq.) dropwise at 0 °C. The reaction mixture was stirred for 1 h at room temperature. The mixture was diluted with DCM (30 mL) and washed with water, HCl 1M and brine. The extracted organic layer was dried over anhydrous $MgSO_4$, filtered and evaporated to afford the desired product as a yellow solid (1.38 g, 97%). $^1H$ NMR is in accordance with the literature[3].

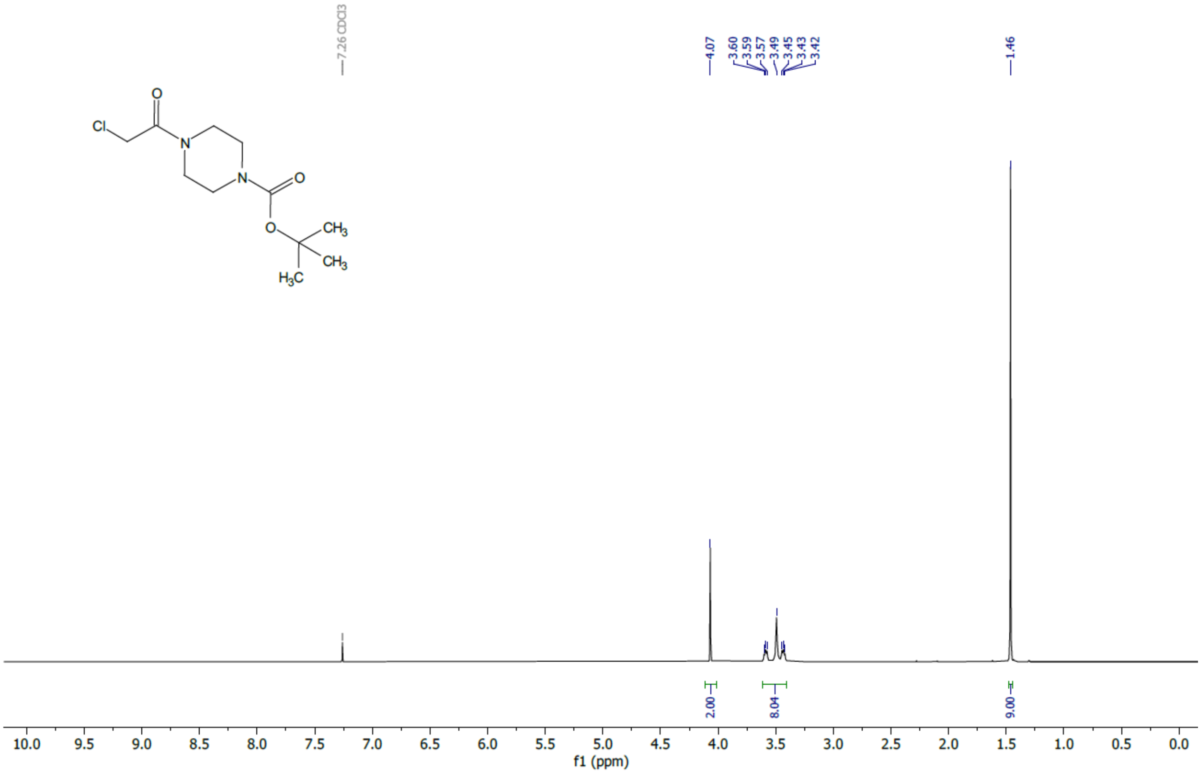


$^1H$ NMR (400 MHz, $CDCl_3$) δ 4.07 (s, 2H), 3.61 – 3.40 (m, 8H), 1.46 (s, 9H).

$^1H$ NMR spectrum of **1**

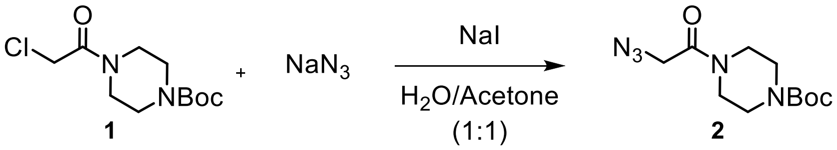


**2.** Piperazine **1** (1.4 g, 5.5 mmol, 1 eq.) was dissolved in a $H_2O$/Acetone mixture (1:1, 20 mL). NaI (0.99 g, 6.6 mmol, 1.2 eq.) was added and reaction mixture stirred for 5 - 10 minutes. $NaN_3$ (0.72 g, 11 mmol, 2 eq.) was added portion wise and reaction mixture stirred at 80°C overnight. Acetone was evaporated under vaccum. Water was added (50 mL) and the product was extracted with DCM (3 x 50 mL). Combined organic layers were dried over anhydrous $MgSO_4$, filtered and evaporated to afford the desired product as a white solid (1.31 g, 88%). $^1H$ NMR is in accordance with the literature[4] .

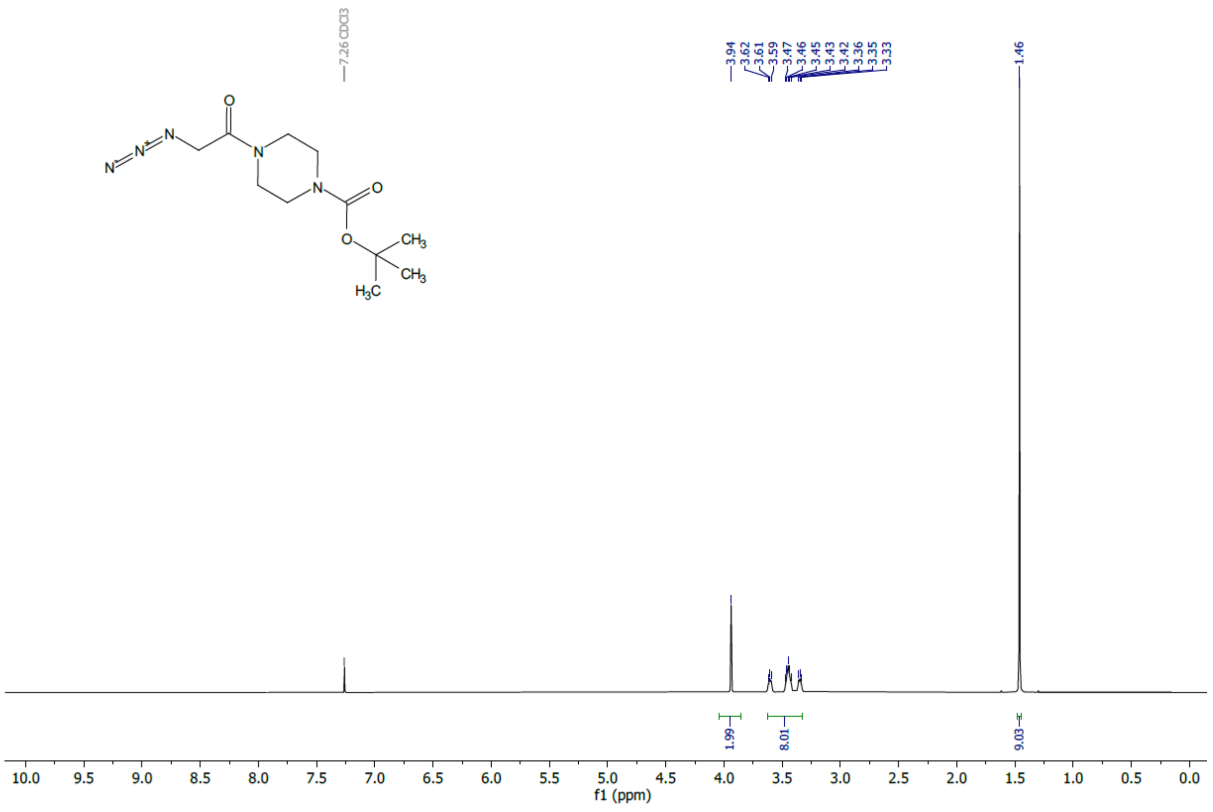


$^1$H NMR (400 MHz, $CDCl_3$) δ 3.94 (s, 2H), 3.63 – 3.33 (m, 8H), 1.46 (s, 9H).

$^1$H NMR spectrum of **2**

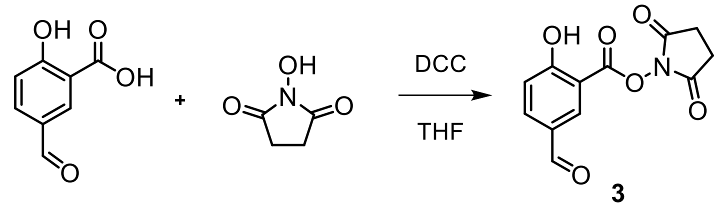


**3**

**3.** To a suspension of 5-formyl salicylic acid (1 g, 6.0 mmol, 1 eq.) and N-hydroxysuccinimide (0.76 g, 6.6 mmol, 1.1 eq.) in THF (10 mL) was added DCC (1.37 g, 6.6 mmol, 1.1 eq.). The mixture was stirred at room temperature for 1 hour and filtered off over a pad of celite. The filtrate was concentrated under reduced pressure then the residue was purified by flash chromatography (heptane/ethyl acetate 7:3 to 3:7) to give **3** (1.1 g, 69 %) as a white solid. $^1$H NMR is in accordance with the literature[5].

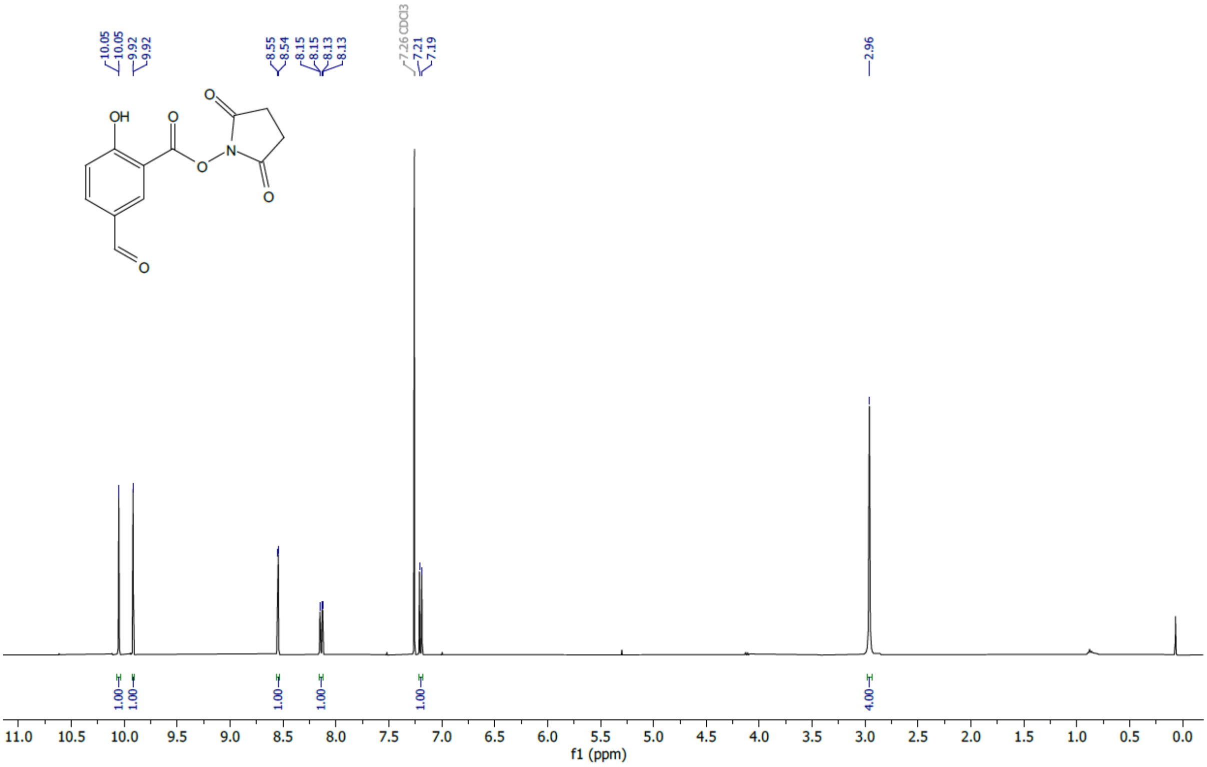


$^1$H NMR (400 MHz, $CDCl_3$) δ 10.05 (d, *J* = 0.5 Hz, 1H), 9.92 (d, *J* = 0.6 Hz, 1H), 8.55 (d, *J* = 2.1 Hz, 1H), 8.14 (dd, *J* = 9.0, 1.9 Hz, 1H), 7.20 (d, *J* = 8.8 Hz, 1H), 2.96 (s, 4H).

$^1$H NMR spectrum of **3**

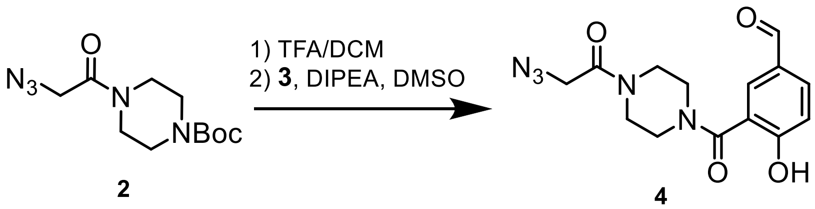


**2** **4**

**4. 2** (0.32 g, 1.2 mmol, 1.2 eq.) was deprotected by dissolving it into a mixture of TFA/DCM (1:1, 5 mL) for 2 hours at room temperature. DCM and TFA were removed *in vacuo*. Deprotected product was added into a solution of **3** (0.26 g, 1.0 mmol, 1 eq.) in 10 mL DMSO, followed by DIPEA (0.5 mL, 3.0 mmol, 3 eq.). The solution was stirred overnight at room temperature. Crude mixture was poured into HCl 1M (10 mL) and extracted with ethyl acetate (3 x 10 mL). The combined organic layers were dried over anhydrous $MgSO_4$, filtered and concentrated. The product was purified by flash chromatography (DCM/ethyl acetate 7:3 to 3:7) to give **4** (153 mg, 49%) as a white solid. Rf = 0.3 (DCM/ethyl acetate 4:6).

$^1$H NMR (400 MHz, $CDCl_3$) δ 9.87 (s, 1H, O=C-<u>H</u>), 7.89 (dd, *J* = 8.5, 2.1 Hz, 1H, $H_{ar}$), 7.83 (d, *J* = 2.1 Hz, 1H, $H_{ar}$), 7.16 (d, *J* = 8.5 Hz, 1H, $H_{ar}$), 4.00 (s, 2H, C$\underline{H_2}$-$N_3$), 3.86 – 3.71 (m, 6H, $H_{piperazine}$), 3.56 – 3.49 (m, 2H, $H_{piperazine}$).

$^{13}$C NMR (400 MHz, $CDCl_3$+MeOD-*d4*) δ 190.59 (O=<u>C</u>-H), 167.98 (O=<u>C</u>-N), 166.41 (O=<u>C</u>-N), 159.73 ($\underline{C_{ar}}$-OH), 133.00 ($C_{ar}$), 131.25 ($C_{ar}$), 128.80 ($C_{ar}$), 122.76 ($C_{ar}$), 116.58 ($C_{ar}$), 50.68 (C-$N_3$), 44.87 ($C_{piperazine}$), 41.93 ($C_{piperazine}$).

HRMS ($ESI^+$): *m/z* calculated for $C_{14}H_{15}N_5O_4$: 317.1124 [M+H]+, found 318.1202.

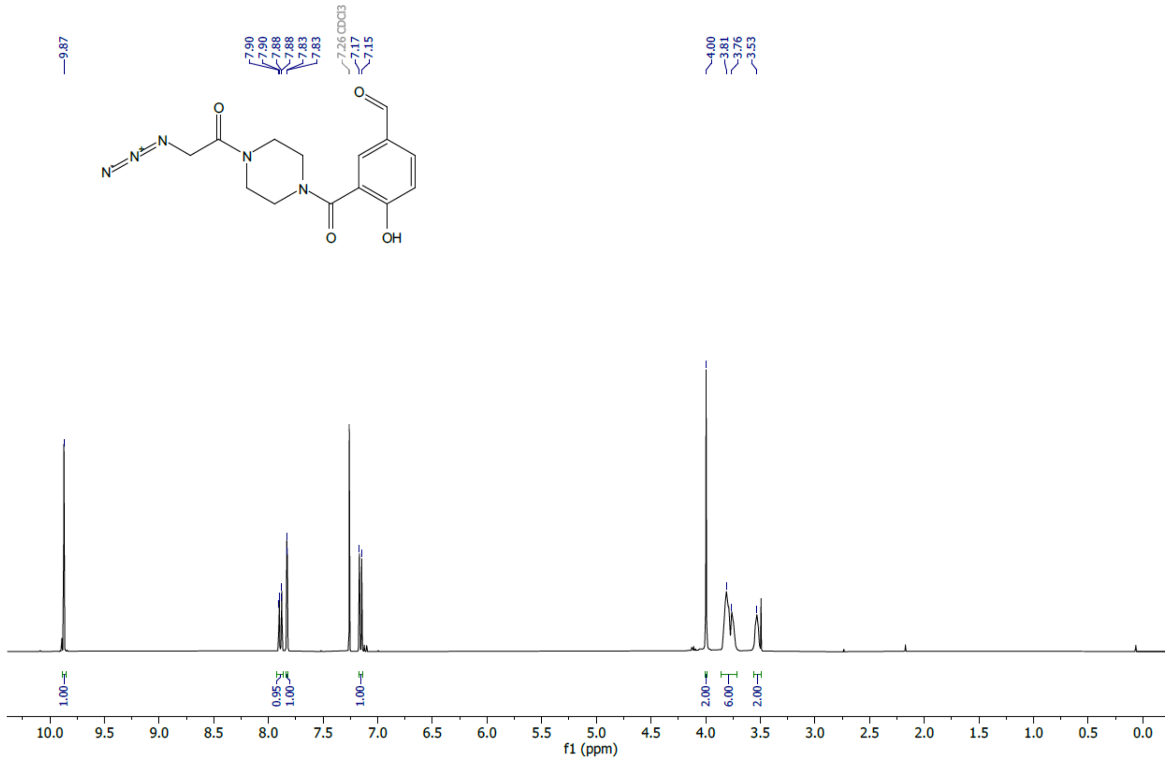

[1]H NMR (400 MHz, $CDCl_3$) δ 9.87 (s, 1H), 7.89 (dd, $J$ = 8.5, 2.1 Hz, 1H), 7.83 (d, $J$ = 2.1 Hz, 1H), 7.16 (d, $J$ = 8.5 Hz, 1H), 4.00 (s, 2H), 3.86 – 3.71 (m, 6H), 3.56 – 3.49 (m, 2H).

$^{1}$H NMR spectrum of **4**

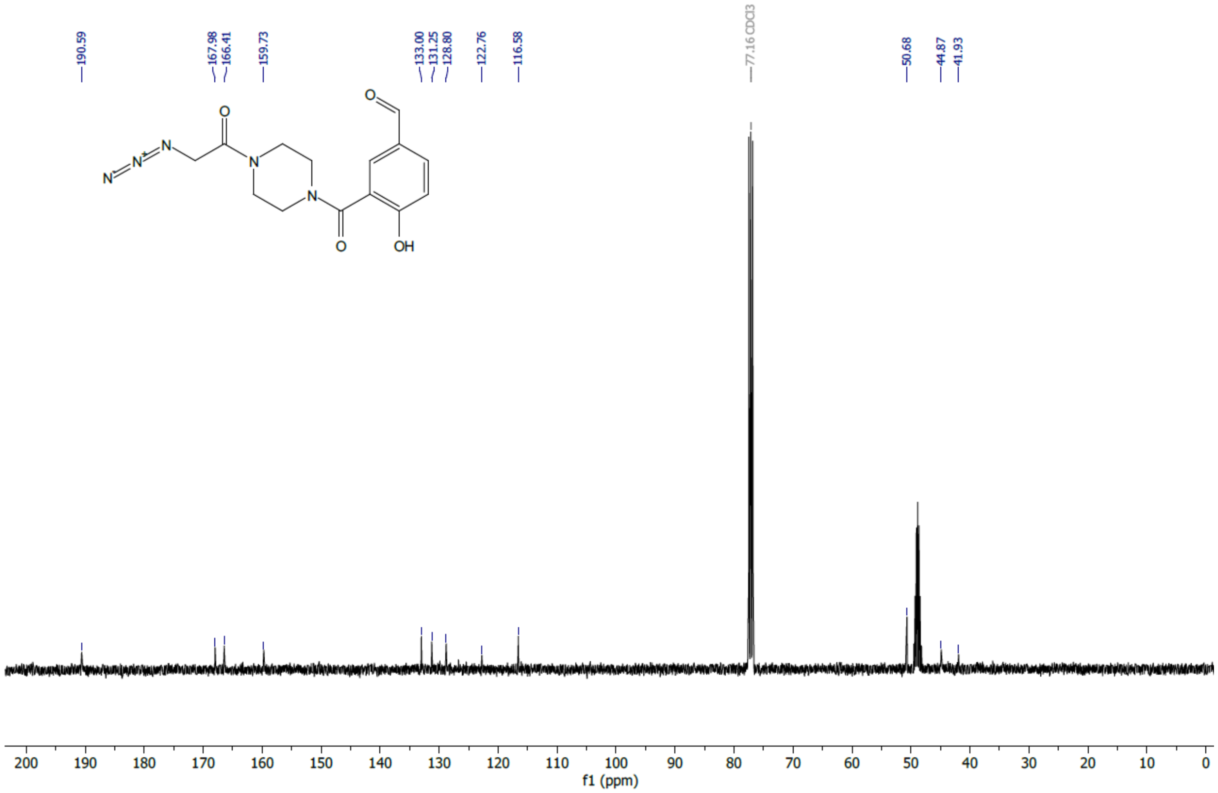

[13]C NMR (400 MHz, $CDCl_3$+MeOD-$d_4$) δ 190.59, 167.98, 166.41, 159.73, 133.00, 131.25, 128.80, 122.76, 116.58, 50.68, 44.87, 41.93.

$^{13}$C NMR spectrum of **4**

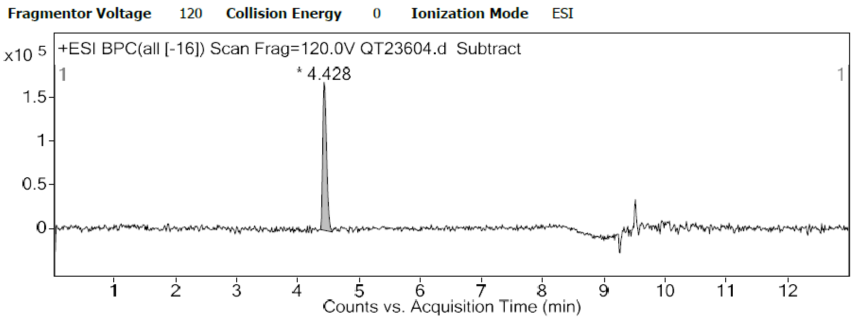


HPLC trace of **4**

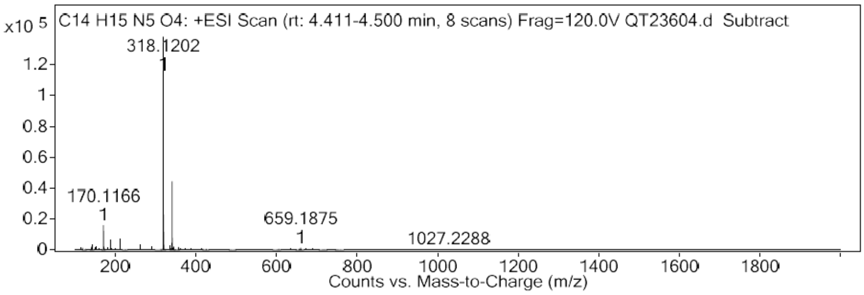


HRMS Spectrum of **4**

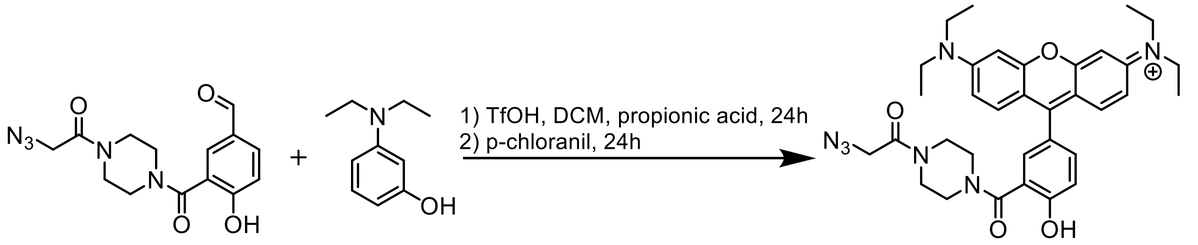


**4** **5**

**5 (H-Ruby-N$_3$). 4** (350 mg, 1.1 mmol, 1 eq.) and 3-diethylaminophenol (401 mg, 2.4 mmol, 2.2 eq.) were dissolved in DCM (5 mL). Triflic acid was added (29 µL, 0.3 mmol, 0.3 eq.), followed by propionic acid (2 mL) to help with solubility. Reaction mixture was stirred in the dark at room temperature for 24 h, until the starting material was completely converted into the aldehyde condensation product. Then chloranil (271 mg, 1.1 mmol, 1 eq.) was added and the reaction mixture was stirred for 24 h in the dark at room temperature. Solvents were evaporated *in vacuo* and the residue was purified by flash chromatography (DCM/MeOH 98:2 to 90:10) to give **5** (117 mg, 16%) as a pink solid. Rf = 0.16 (DCM/MeOH 95:5).

Product was not completely soluble in MeOD-*d4* and was not stable in a $CDCl_3$/MeOD-*d4* mixture to obtain a proper NMR analysis.

HRMS (ESI$^+$): *m/z* calculated for $C_{34}H_{40}N_7O_4$: 610.3142 [M]+, found 610.3152.

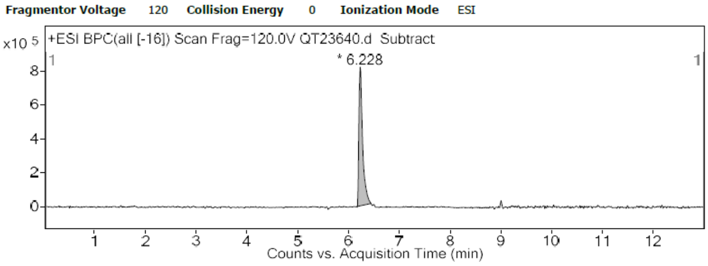


HPLC trace of **5**

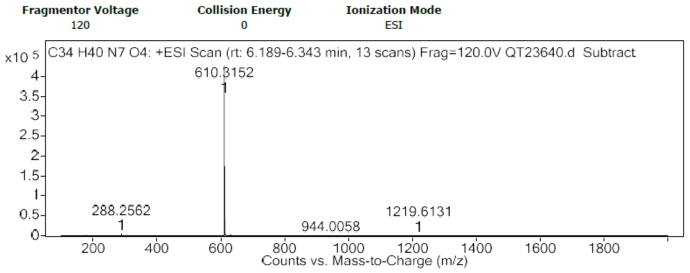


HRMS Spectrum of **5**

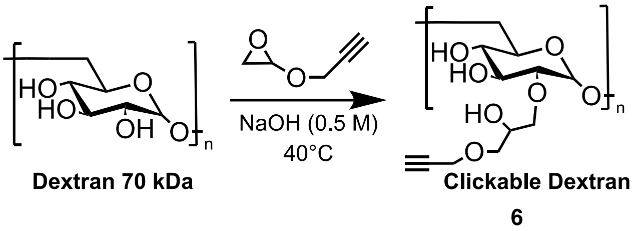


**6. Clickable Dextran.** Same procedure as Despras et al. was followed to obtain 220 mg of Clickable 70 kDa Dextran (Degree of Substitution = 70%, Final MW = 104 kDa).[3]

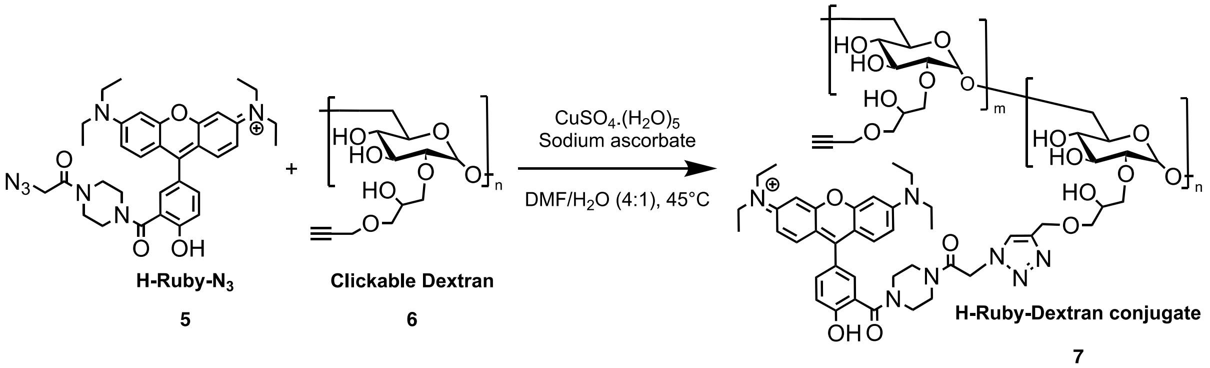


**7. H-Ruby Dextran conjugate.** To a solution of **H-Ruby-N₃** (2.5 mg, 4.1 µmol) and **clickable dextran** (30 mg, 0.3 µmol) in 400 µL degassed DMF was added copper sulfate pentahydrate (5 mg) and sodium ascorbate (5 mg) in water (100 µL). The reaction was stirred in the dark at 40°C for 30 min. DMF was evaporated and the crude residue was dissolved in aq. 0.1 M EDTA (1 mL) then purified over G-25 size exclusion column and lyophilized to obtain **H-Ruby Dextran conjugate** as a pink solid (28 mg, molar ratio ~ 3 mol dye/mol dextran).

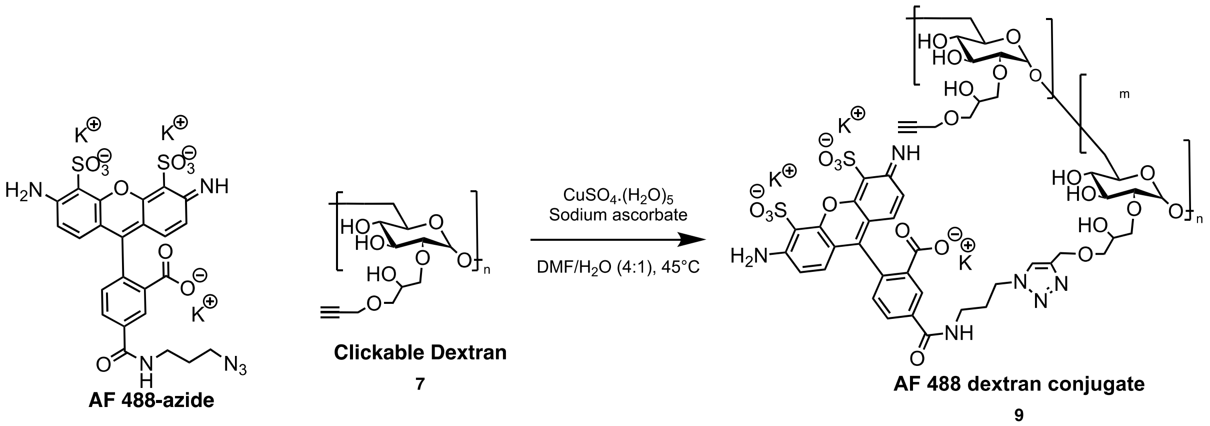


**9. AF488 Dextran conjugate 8.** To a solution of AF488-azide (purchased from Lumiprobe, 2.5 mg, 3.6 µmol) and **clickable dextran** (50 mg, 0.5 µmol) in $H_2O$/DMF (4:1, 5 mL) was added copper sulfate pentahydrate (5 mg) and sodium ascorbate (5 mg) in water (100 µL).

The reaction was stirred in the dark at 45°C for 4 hours. DMF was evaporated and the crude residue was dissolved in aq. 0.1 M EDTA (1 mL) then purified over G-25 size exclusion column and lyophilized to obtain **AF 488 dextran conjugate** as an orange solid (49 mg, molar ratio ~ 1.7 mol dye/mol dextran).
